# High-surety isothermal amplification and detection of SARS-CoV-2, including with crude enzymes

**DOI:** 10.1101/2020.04.13.039941

**Authors:** Sanchita Bhadra, Timothy E. Riedel, Simren Lakhotia, Nicholas D. Tran, Andrew D. Ellington

## Abstract

Isothermal nucleic acid amplification tests (iNAT), such as loop-mediated isothermal amplification (LAMP), are good alternatives to polymerase chain reaction (PCR)-based amplification assays, especially for point-of-care and low resource use, in part because they can be carried out with relatively simple instrumentation. However, iNATs can generate spurious amplicons, especially in the absence of target sequences, resulting in false positive results. This is especially true if signals are based on non-sequence-specific probes, such as intercalating dyes or pH changes. In addition, pathogens often prove to be moving, evolving targets, and can accumulate mutations that will lead to inefficient primer binding and thus false negative results. Internally redundant assays targeting different regions of the target sequence can help to reduce such false negatives. Here we describe rapid conversion of three previously described SARS-CoV-2 LAMP assays that relied on non-sequence-specific readout into assays that can be visually read using sequence-specific fluorogenic oligonucleotide strand exchange (OSD) probes. We evaluate one-pot operation of both individual and multiplex LAMP-OSD assays and demonstrate detection of SARS-CoV-2 virions in crude human saliva.

## INTRODUCTION

Loop-mediated isothermal amplification (LAMP) uses the strand-displacing Bst DNA polymerase and 4 primers (FIP, BIP, F3, and B3) that bind to 6 target regions (B3, B2, B1, F1c, F2c and F3c) to generate 10^9^ to 10^10^ copies of DNA or RNA targets, typically within 1 to 2 h (**Figure 1**).^*1*^ In greater detail, F2 in FIP (F1c-F2) and B2 in BIP (B1c-B2) initiate amplification. F1c and B1c self-prime subsequent amplification. F3 and B3-initiated DNA synthesis displaces FIP and BIP-initiated strands. 3’-ends of the resulting single-stranded, dumbbell-shaped amplicons are extended to hairpins by *Bst* polymerase. FIP and BIP hybridize to the single-stranded loops and initiate DNA synthesis that opens the hairpin to form concatameric amplicons containing self-priming 3’-end hairpins. The ensuing continuous amplification generates double-stranded concatameric amplicons with self-priming hairpins and single-stranded loops.^*1*^

**Figure 1.**
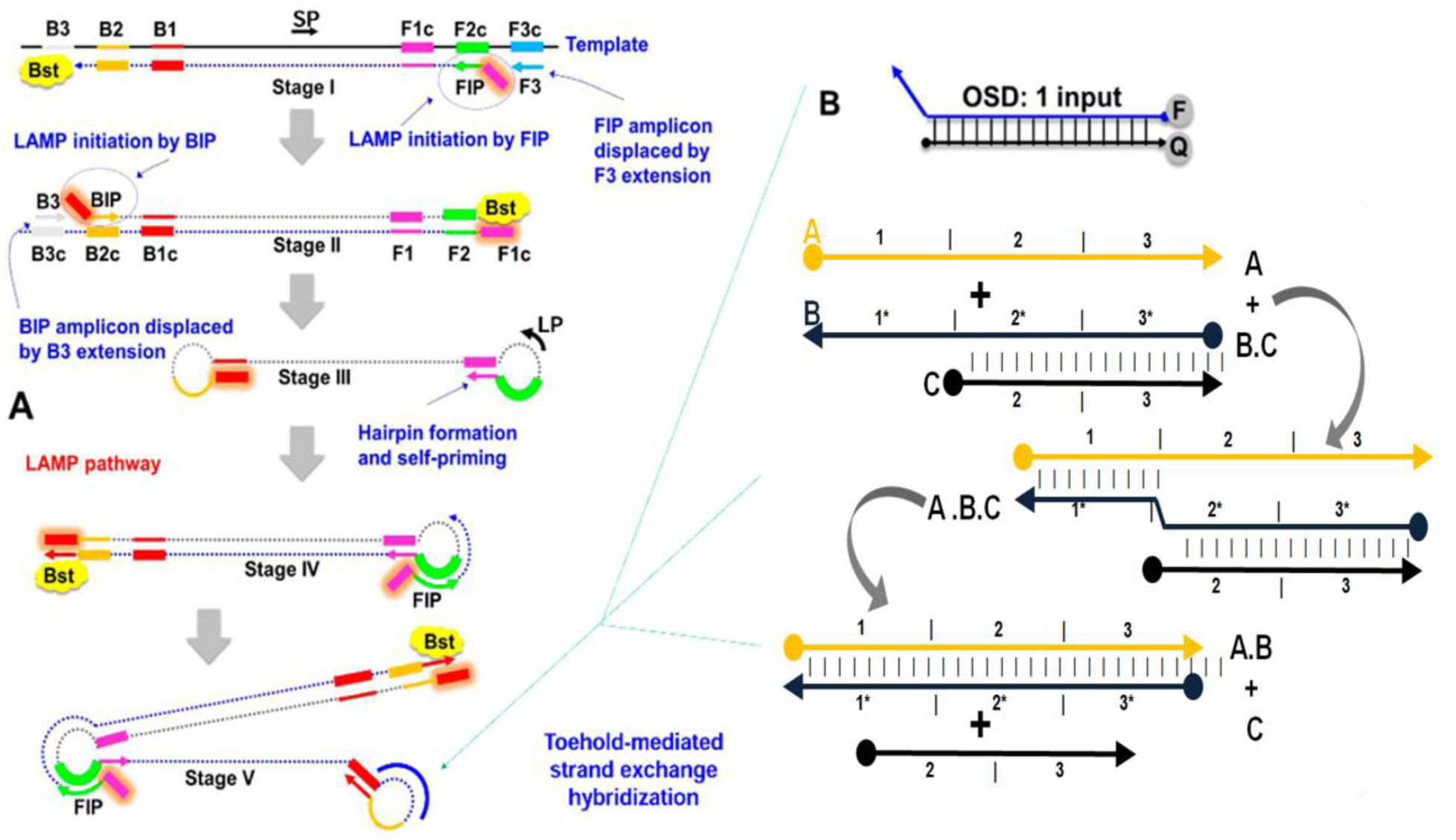
LAMP-OSD schematic. ‘c’ denotes complementary sequences. F and Q on the OSD denote fluorophore and quencher, respectively. OSD and subsequent strand exchange intermediates are denoted by numbered domains, which represent short (usually <12 nt) sequences in an otherwise continuous oligonucleotide. Complementary domains are indicated by asterisk.

LAMP can rival PCR for sensitivity without thermocycling,^*2*^ and additional stem and loop primers can accelerate amplification, with some LAMP assays being complete within 10 min.^*3*, *4*^ However, since LAMP is commonly read using non-specific methods (such as, Mg^2+^ precipitation, intercalating dyes or labeled primers) that cannot distinguish spurious amplicons that frequently arise from continuous amplification, its utility can be limited. We have previously overcome these drawbacks using oligonucleotide strand exchange (OSD) probes, based in part on advances in strand exchange DNA computation (**Figure 1**). Strand exchange occurs when two partially or fully complementary strands hybridize to each other by displacing pre-hybridized strand(s) (**Figure 1B**). Strand exchange usually initiates by basepairing at single-stranded ‘toeholds’ and progresses to form additional basepairs via branch migration, allowing the rational design of complex algorithms and programmable nanostructures^*5*–*9*^. The hemiduplex oligonucleotide probes contain a so-called ‘toehold’ that allows sequence-specific interaction with a target molecule, and have opposed fluor and quencher moieties. In the presence of a complementary target, the OSD probes can undergo strand exchange and separation, leading to an easily read fluorescent signal.^*10*^ In essence the OSD probes are functional equivalents of TaqMan probes and have been shown to accurately report single or multiplex LAMP amplicons from few tens of targets without interference from non-specific amplicons or inhibitors.^*10*, *11*^ Of equal import, the programmability of OSD probes allows their adaptation to many different assay formats, including to off-the-shelf devices such as glucometers and pregnancy test strips.^*12*–*16*^

LAMP-OSD is designed consciously to be easy to use and interpret, which makes it a reliable choice for either screening or validation of disease states. Base-pairing to the toehold region is extremely sensitive to mismatches, ensuring specificity, and the programmability of both primers and probes makes possible rapid adaptation to new diseases or new disease variants. We have shown that higher order molecular information processing is also possible, such as integration of signals from multiple amplicons^*17*^. Overall, the use of sequence-specific probes allow construction of strand exchange computation circuits that act as ‘matter computers’,^*5*–*8*^ something that is not generally possible within the context of a PCR reaction (which would of necessity melt the computational devices).

We have taken pains to make LAMP-OSD robust for resource-poor settings. Lyophilized master mixes are stable without cold chain for extended durations and can be operated simply upon rehydration and addition of crude sample.^*18*^ The one-pot operation, direct analysis of crude specimens, and easy yes/no visual readout make LAMP-OSD ideal for field operation with minimal training and resources.

OSD probes can be readily designed for integration into existing LAMP assays without significant disruption to standard assay practice. To that end, here we demonstrate the conversion of three recently pre-published LAMP primer sets described for detection of SARS-CoV-2, but that used non-specific readout methods (**Table 1**). The LAMP-OSD versions of these assays maintain the simplicity of visual yes/no readout, while endowing the assays with the inherent accuracy of probe-based signal transduction. We also perform multiplex execution of some of these LAMP-OSD assays and demonstrate the feasibility of sample-to-answer operation by directly analyzing human saliva spiked with SARS-CoV-2 virions.

**Table 1.**
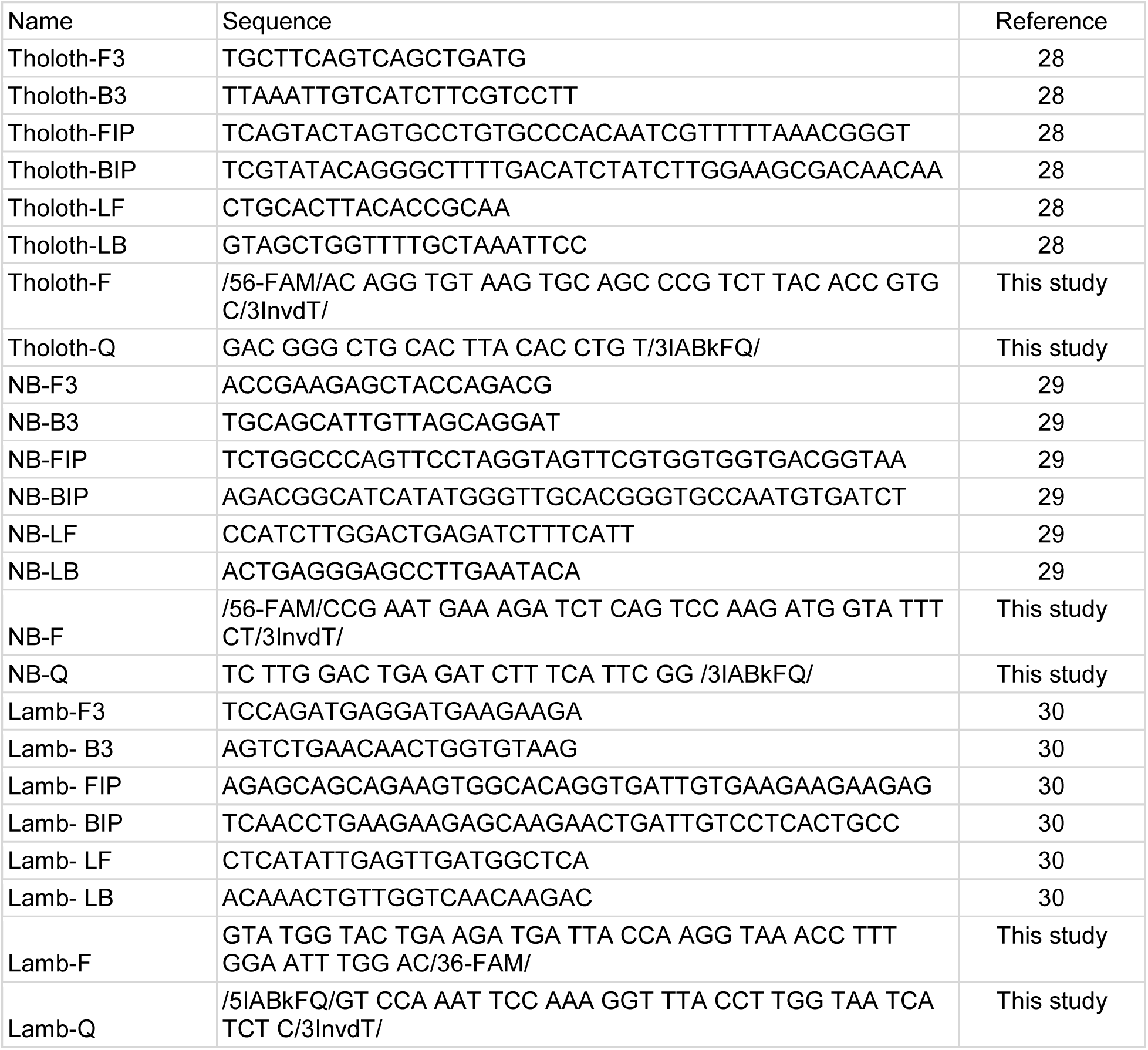
LAMP primers and OSD probes.

## METHODS

### Chemicals and reagents

All chemicals were of analytical grade and were purchased from Sigma-Aldrich (St. Louis, MO, USA) unless otherwise indicated. All enzymes and related buffers were purchased from New England Biolabs (NEB, Ipswich, MA, USA) unless otherwise indicated. All oligonucleotides and gene blocks (**Table 1**) were obtained from Integrated DNA Technologies (IDT, Coralville, IA, USA). SARS-CoV-2 N gene synthetic transcript was a gift from the Schoggins lab at UT Southwestern Medical Center, Dallas, TX. SARS-CoV-2 genomic RNA and heat-inactivated virions were obtained from American Type Culture Collection.

### OSD probe design

We chose to design OSD probes for three of these pre-published LAMP primer sets, from here on referred to as the Tholoth, Lamb, and NB primers. The three primer sets target different regions in the ORF1AB and N genes of the SARS-CoV-2 genome. Fluorogenic OSD probes were designed for each of these primer sets using previously described principles and the Nucleic acid circuit design software NUPACK available freely at http://www.nupack.org/. ^*10*, *19*^ Briefly, the target derived loop regions between the F1c and F2 primer binding sites were chosen as OSD binding regions for each of the three LAMP primer sets. The long OSD strand was designed to be complementary to this loop region. Single stranded 10-12 nucleotides long toehold regions were designated on one end of this long strand while a complementary short OSD strand was designed to hybridize to the remaining portion of the long strand. The long strand was labeled with a fluorescein moiety at the terminus not acting as the toehold. The short strand was labeled with a quencher and all free 3’-OH ends were blocked with inverted dT to prevent extension by DNA polymerase.

### Reverse transcription (RT) LAMP assay

Individual LAMP assays were assembled in a total volume of 25 μl of 1X Isothermal buffer (NEB; 20 mM Tris-HCl, 10 mM (NH_4_)_2_SO_4_, 50 mM KCl, 2 mM MgSO_4_, 0.1% Tween 20, pH 8.8 at 25°C). The buffer was supplemented with 1.4 mM dNTPs, 0.4 M betaine, 6 mM additional MgCl_2_, 2.4 μM each of FIP and BIP, 1.2 μM of indicated loop primers, 0.6 μM each of F3 and B3 primers, 16 units of *Bst* 2.0 DNA polymerase, and 7.5 units of warmstart RTX reverse transcriptase. Amplicon accumulation was measured by adding OSD probes. First, Tholoth, Lamb, and NB OSD probes were prepared by annealing 1 μM of the fluorophore-labeled OSD strand with 2 μM, 3 μM, and 5 μM, respectively of the quencher-labeled strand in 1X Isothermal buffer. Annealing was performed by denaturing the oligonucleotide mix at 95 °C for 1 min followed by slow cooling at the rate of 0.1 °C/s to 25 °C. Excess annealed probes were stored at −20 °C. Annealed Tholoth, Lamb, and NB OSD probes were added to their respective LAMP reactions at a final concentration of 100 nM, 100 nM, and 120 nM, respectively of the fluorophore-bearing strand. Multiplex LAMP-OSD assays comprising both Tholoth and NB primers and probes were set up using the same conditions as above except, the total LAMP primer amounts were made up of equimolar amounts of Tholoth and NB primers.

In some experiments, LAMP-OSD reactions were assembled using Bst-LF instead of Bst 2.0 and RTX. This was done by replacing Bst 2.0 and RTX in the LAMP-OSD reactions with ~2 x 10^7^ CFU of rehydrated Bst-LF cellular reagents.^*20*^ These cellular reagents were prepared in an *E. coli* BL21(DE3) derivative ΔendA ΔrecA strain using previously described protocols.^*20*^ Briefly, chemically competent bacteria were transformed with Bst-LF expression plasmid. Overnight 3 ml cultures of these transformed bacteria were grown in 2X YT broth containing 100 μg/mL ampicillin. Subsequently, sub-cultures were initiated at 1:200 dilution using 50 mL Superior Broth™ (Athena Environmental Sciences, Inc., Baltimore, MD, USA) containing 100 μg/ml ampicillin. These cultures were incubated at 37 °C and constant 225 rpm agitation till they reached the log phase of growth (A600 = 0.4 to 0.7). Protein production was initiated by adding 20ng/ml anhydrotetracycline (aTC) followed by 3 h incubation at 37 °C with 225 rpm agitation.

After induction, bacteria were collected by centrifugation followed by washing twice in cold 1X PBS (137 mM NaCl, 2.7 mM KCl, 4.3 mM Na_2_HPO_4_, 1.47 mM KH_2_PO_4_, pH 7.4). The bacterial pellets were resuspended in cold 1X PBS at a density of A_600_ = 3.5 to 6.5. Some 2×10^8^ induced bacteria (estimated from the A_600_ value using the relation 0.5 optical density = 5×10^8^ bacteria/ml) were aliquoted into individual 0.2 ml PCR tubes and frozen at −80 °C overnight prior to lyophilization for 3 h at 197 mTorr and −108 °C using a VirTis Benchtop Pro lyophilizer (SP Scientific, Warminster, PA, USA). Immediately before use, each tube of cellular reagents was hydrated in 30 μL water and 3 μL of this suspension was added to each LAMP reaction.

Templates were serially diluted in TE buffer (10 mM Tris-HCl, pH 7.5:0.1 mM EDTA, pH 8.0) immediately prior to use. In some experiments, templates were diluted in human saliva that had been heated at 95 °C for 10 min. Templates used included: zero to several hundred copies of synthetic double stranded linear DNA gBlock (IDT, Coralville, Iowa, USA), *in vitro* transcribed RNA, viral genomic RNA, and heat-inactivated virions. Following addition of templates to RT-LAMP-OSD reagents, reaction mixes were incubated at 65 °C for 90 min.

Some LAMP-OSD reactions were analyzed in real-time using LightCycler 96 real-time PCR machine (Roche, Basel, Switzerland). Reactions were subjected to 30 cycles of two-step incubations – step 1:150 sec at 65 °C, step 2: 30 sec at 65 °C. Fluorescence was measured in the FAM channel during step 2 of each cycle. LAMP-OSD assays intended for visual ‘yes/no’ readout of endpoint fluorescence were assembled in 0.2 ml optically clear thin-walled tubes with low auto-fluorescence (Axygen, Union City, CA, USA). Following 90 min of amplification at 65 °C, endpoint fluorescence was imaged using either a cellphone and a blue LED transilluminator or a BioRad ChemiDoc camera (Bio-Rad Laboratories, Hercules, CA, USA).

## RESULTS

### Integration of OSD probes into pre-published SARS-CoV-2 LAMP primer sets

A series of 11 published primer sets for SARS-CoV-2 were screened using New England Biolab’s WarmStart^®^ Colorimetric LAMP 2X Master Mix (NEB, Ipswich, MA, USA) according to the manufacturer’s protocol (**Table 2**). We found that 9 of the 11 sets showed significant no-template amplification in over 10% of the replicates in less than an hour of incubation at 65 °C (data not shown). These results are consistent with other published results that rely on colorimetric LAMP reactions, rather than on probe-based detection.^*21*^ In fact, for many published assays, color changes must be read within a narrow window of time in order to minimize spurious conclusions, a consideration that does not scale well for diagnostic screening, especially at point-of-care or as an early part of a clinical diagnostics pipeline.

**Table 2.** Pre-published LAMP primer sets for SARS-CoV-2 found online before March 04, 2020.

| Reference | Preprint Date | Gene Target | Temp | Validation |
| --- | --- | --- | --- | --- |
| 28 | 2020.02.16 | ORF1ab | 63 | Synthesized DNA |
| 30 | 2020.02.19 | ORF1a | 65 | Synthetic Patient Samples |
| 31 | 2020.02.20 | ORF1ab | 63 | Patient Samples |
| 29 | 2020.02.26 | ORF1a-A | 65 | Patient samples |
| 29 | 2020.02.26 | ORF1a-B | 65 | Patient samples |
| 29 | 2020.02.26 | ORF1a-C | 65 | Patient samples |
| 29 | 2020.02.26 | Gene N-A | 65 | Patient samples |
| 29 | 2020.02.26 | Gene N-B | 65 | Patient samples |
| 32 | 2020.03.02 | ORF1ab | 63 | Patient samples |
| 32 | 2020.03.02 | N | 63 | Patient samples |
| 32 | 2020.03.02 | E | 63 | Patient samples |

To suppress potential false positive readout, we chose to develop OSD probes for three of the LAMP primer sets, termed herein as NB, Lamb, and Tholoth (**Table 1**). These primer sets target three different regions of the viral genome, the N gene, the NSP3 coding region of ORF1AB, and the RNA-dependent RNA polymerase coding region of ORF1AB. Of the three primer sets, the NB assay had the lowest propensity for spurious signal when analyzed by non-specific colorimetric readout (data not shown). To create LAMP-OSD versions of these assays, we designed OSD probes that were complementary to one of the loop sequences in each of the three LAMP amplicons. Subsequently, Tholoth, Lamb, and NB LAMP-OSD assays were setup individually by mixing separate reaction components as indicated in the **Methods** section. Each individual assay contained its specific OSD probes along with both inner primers FIP and BIP and both outer primers F3 and B3. In addition, each assay also received the backward loop (LB) primer that bound to the amplicon loop between B1c and B2 sites that was not recognized by the respective OSD probe. The forward loop (LF) primer that overlapped the Tholoth and NB OSD binding region was excluded. The LF primer was also initially excluded in Lamb LAMP-OSD assay even though the amplicon loop that bound this loop primer was long enough to accommodate a non-overlapping OSD reporter; this was done to fairly compare the amplification kinetics of all three assays in a 5-primer format. In later versions of the assay with the Lamb primers, all 6 primers were included (designated as “6 primer Lamb” in **Figures 3**, **6**, and **7**).

As shown in **Figure 2**, in response to target templates, all three LAMP-OSD assays generated strong OSD signal that could be measured both in real-time and observed visually at endpoint without interference from noise.

**Figure 2.**
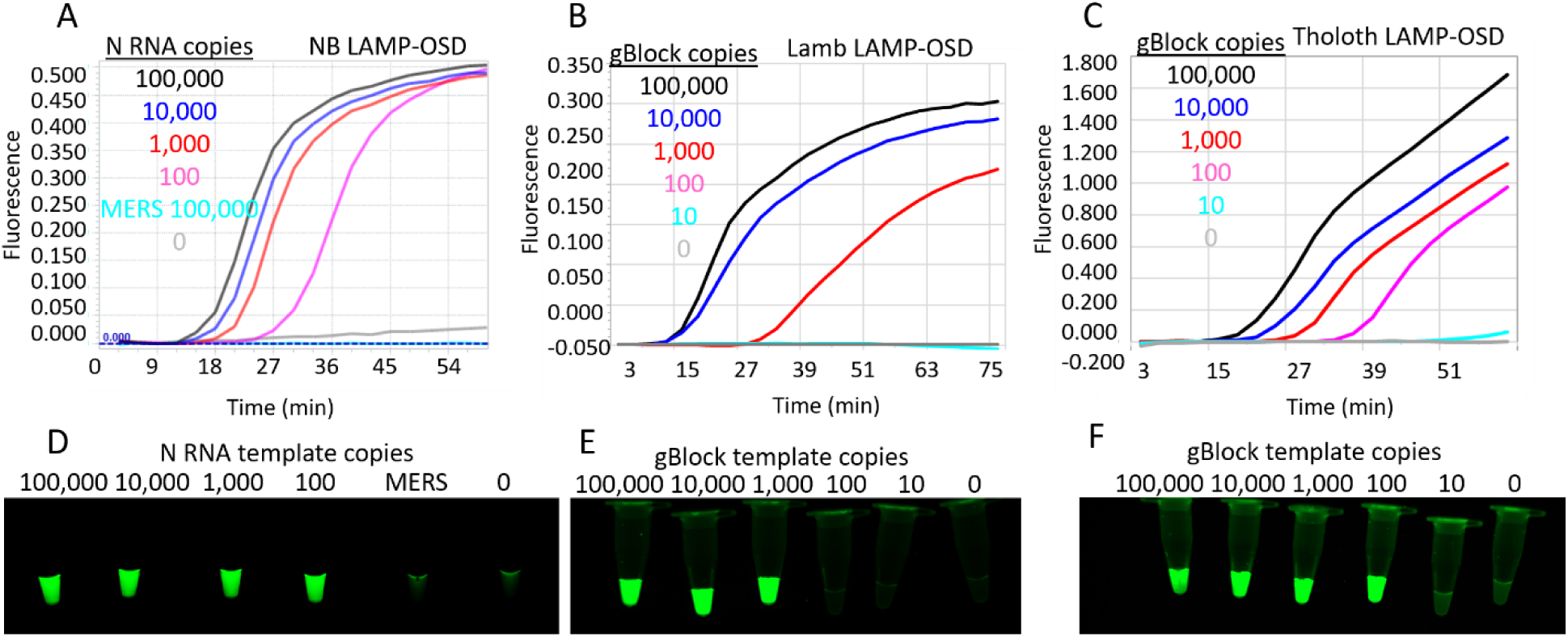
SARS-CoV-2 LAMP-OSD assays. OSD fluorescence measured in real-time during LAMP amplification for NB (A), 5 primer Lamb (B), and Tholoth (C) LAMP-OSD assays are depicted as amplification curves. Presence or absence of OSD fluorescence visually observed at assay endpoint for NB (D), Lamb (E), and Tholoth (F) LAMP-OSD assays are depicted as images of reaction tubes. NB LAMP-OSD assays were seeded with indicated copies of SARS-CoV-2 N RNA or MERS-CoV N RNA or no templates. Lamb and Tholoth LAMP-OSD assays were seeded with indicated copies of gBlock DNA templates.

We then tested the three LAMP-OSD assays using SARS-CoV-2 genomic RNA as templates. While the NB and Tholoth LAMP-OSD assays were performed using 5 primers (FIP, BIP, F3, B3, and LB), the Lamb LAMP-OSD assay was tested using either 5 primers (FIP, BIP, F3, B3, and LB) or 6 primers (FIP, BIP, F3, B3, LB, and LF). Following 90 min of amplification at 65 °C, presence or absence of OSD fluorescence at endpoint was visually observed. As shown in **Figure 3**, presence of SARS-CoV-2 genomic RNA resulted in bright, easily detected fluorescence in all three LAMP-OSD assays. The 6-primer version of Lamb LAMP-OSD could detect fewer genomic RNA copies compared to the 5-primer version of the assay. In contrast, all assays showed no signal in the presence of only human genomic DNA.

**Figure 3.**
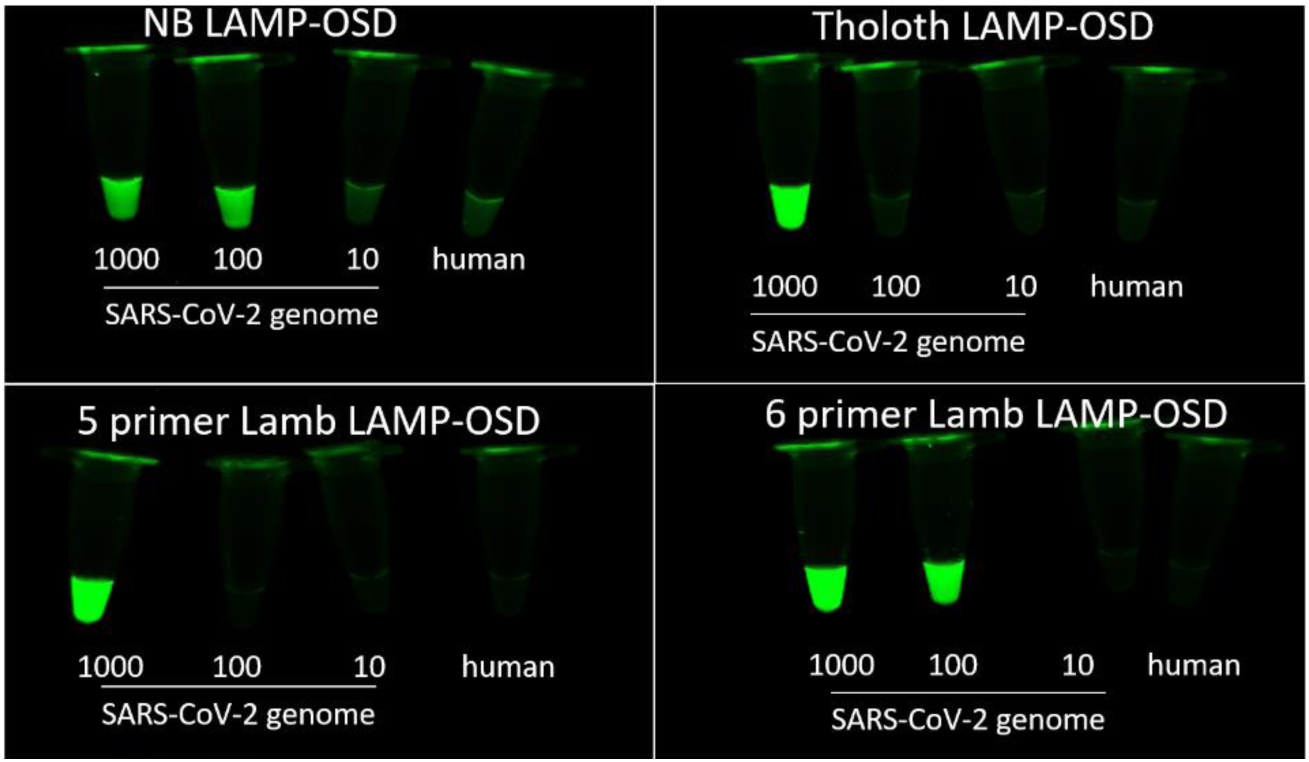
LAMP-OSD analysis of SARS-CoV-2 genomic RNA. Indicated copies of SARS-CoV-2 genomic RNA were analyzed using NB, Tholoth, and both 5 primer and 6 primer Lamb LAMP-OSD assays. Negative control assays received 23 ng of human genomic DNA. Images of endpoint OSD fluorescence are depicted.

### Multiplex LAMP-OSD assay for SARS-CoV-2

Multiplex assays designed to detect multiple sequences from an organism are often employed to improve the accuracy of identification.^*22*, *23*^ CDC recommended diagnostic protocol for SARS-CoV-2 includes RT-qPCR amplification of at least two different regions of the viral genome. In fact, a recent pre-publication demonstrated a multiplex PCR approach to enhance efficiency of detecting SARS-CoV-2 at low copy numbers.^*24*^

Having determined that the individual LAMP-OSD assays with NB, Tholoth, and Lamb primers could signal the presence of SARS-CoV-2 RNA, we sought to execute these assays in a multiplexed format to create an internally redundant assay for SARS-CoV-2. We chose to multiplex the NB and the Tholoth assays because they target different viral genes: the N gene and the ORF1AB gene, respectively. We first tested the ability of both primer sets to amplify their respective synthetic targets (*in vitro* RNA transcripts of N gene and ORF1AB gBlock DNA templates) in a multiplex assay format by assembling LAMP-OSD reactions containing equimolar amounts of both Tholoth and NB LAMP primer sets with either only one or both OSD probes.

When these multiplex assays were seeded with both types of target templates, both Tholoth and NB primer sets led to an increase in OSD fluorescence, measured both in real-time and visually observed at endpoint (**Figure 4**). Multiplex assays containing both OSD probes demonstrated an additive effect, with OSD signal being brighter than assays containing only one type of OSD.

**Figure 4.**
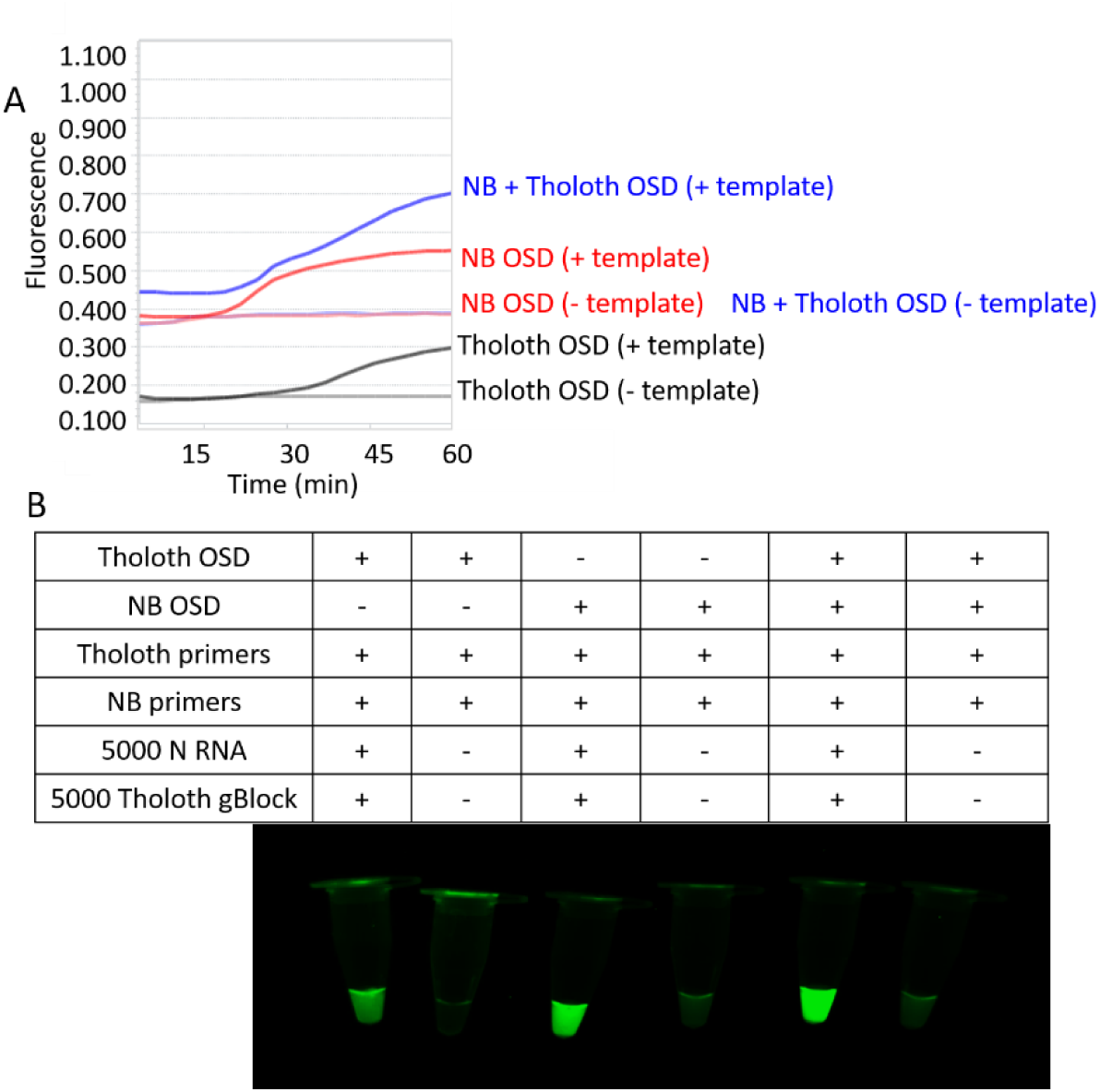
Multiplex LAMP-OSD assay for SARS-CoV-2. Tholoth and NB LAMP-OSD assays were combined in a multiplex format and analyzed using either individual or both OSD probes. (A) Amplification curves showing real-time increase in OSD fluorescence. (B) Images of endpoint OSD fluorescence in multiplex LAMP-OSD assays.

Having confirmed that both the NB and Tholoth primer sets are able to amplify their respective targets in one-pot individual and multiplex reactions containing N gene and ORF1AB sequences (synthetic or viral, **Figures 3**, **4**), we tested the multiplex assay containing both Tholoth and NB primers and OSD probes using full length SARS-CoV-2 viral genomic RNA (**Figure 5**). Visual observation of endpoint fluorescence revealed a bright signal in multiplex assays containing as few as 10 copies of genomic RNA while reactions containing non-specific human DNA remained dark (**Figure 5**). Compared to individual LAMP-OSD assays of SARS-CoV-2 viral genomic RNA (**Figure 3**), it seems that multiplex LAMP-OSD assay is detecting fewer genomic RNA copies (**Figure 5**). For instance, while NB and Tholoth LAMP-OSD assays could routinely detect 100 and 1000 genomic RNA copies, respectively, the multiplex LAMP-OSD assay could signal the presence of as few as 10 genomic RNA copies.

**Figure 5.**
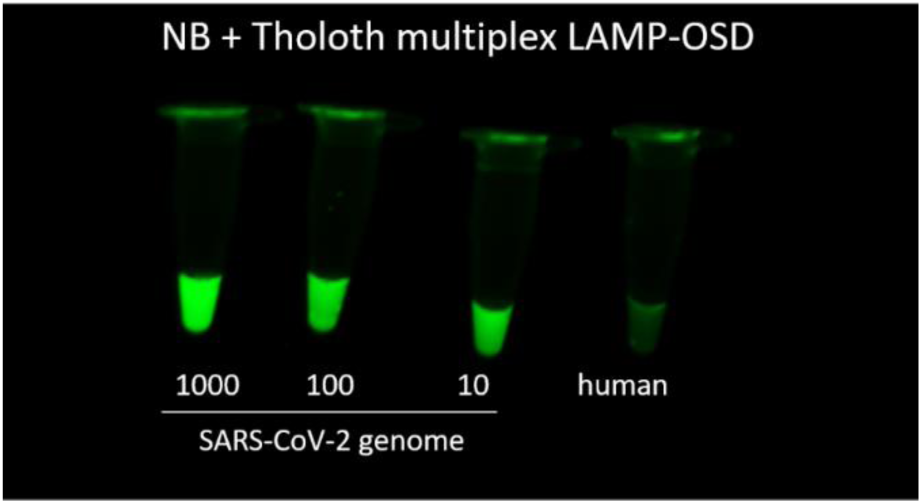
Multiplex LAMP-OSD analysis of SARS-CoV-2 genomic RNA. All assays contained equimolar amount of both Tholoth and NB LAMP primers as well as both Tholoth and NB OSD probes. RNA copies in each reaction are indicated below the respective tube. Control reaction received 23 ng of human genomic DNA. Image of endpoint OSD fluorescence is depicted.

### Direct LAMP-OSD analysis of SARS-CoV-2 virion-spiked human saliva

Given the low limits of detection we have observed, it is possible that LAMP-OSD might be used as part of diagnostics pipelines, or in direct patient screening. However, for this the reactions would need to operate under conditions commensurate with sample collection, especially in resource poor settings. Collection of nasopharyngeal and oropharyngeal swab specimens causes considerable discomfort to patients and requires supplies in the form of sterile swabs and transport media. Moreover, these samples are relatively difficult to self-collect. In contrast, saliva can be non-invasively collected simply by spitting in a sterile collection vessel and it can be done just as easily in a clinic as well as at home. Moreover, studies have shown that SARS-CoV-2 can be consistently detected in patient saliva with median viral loads of 3.3 × 10^6^ copies/mL.^*25*^

We tested the direct sample analysis ability of individual and multiplex LAMP-OSD assays by seeding them with 3 μL of human saliva spiked with SARS-CoV-2 virions. As control, duplicate LAMP-OSD reactions were seeded with virions suspended in 3 μL of TE buffer. Following 90 min incubation at 65 °C, endpoint observation of presence or absence of OSD fluorescence revealed that all assays seeded with SARS-CoV-2 virions, whether in human saliva or TE buffer, were brightly fluorescent (**Figure 6**). In contrast, assays lacking specific templates remained dark. These results suggest that LAMP-OSD assays might be used for direct analysis of human saliva samples in order to amplify and detect genetic signatures from SARS-CoV-2 virions.

**Figure 6.**
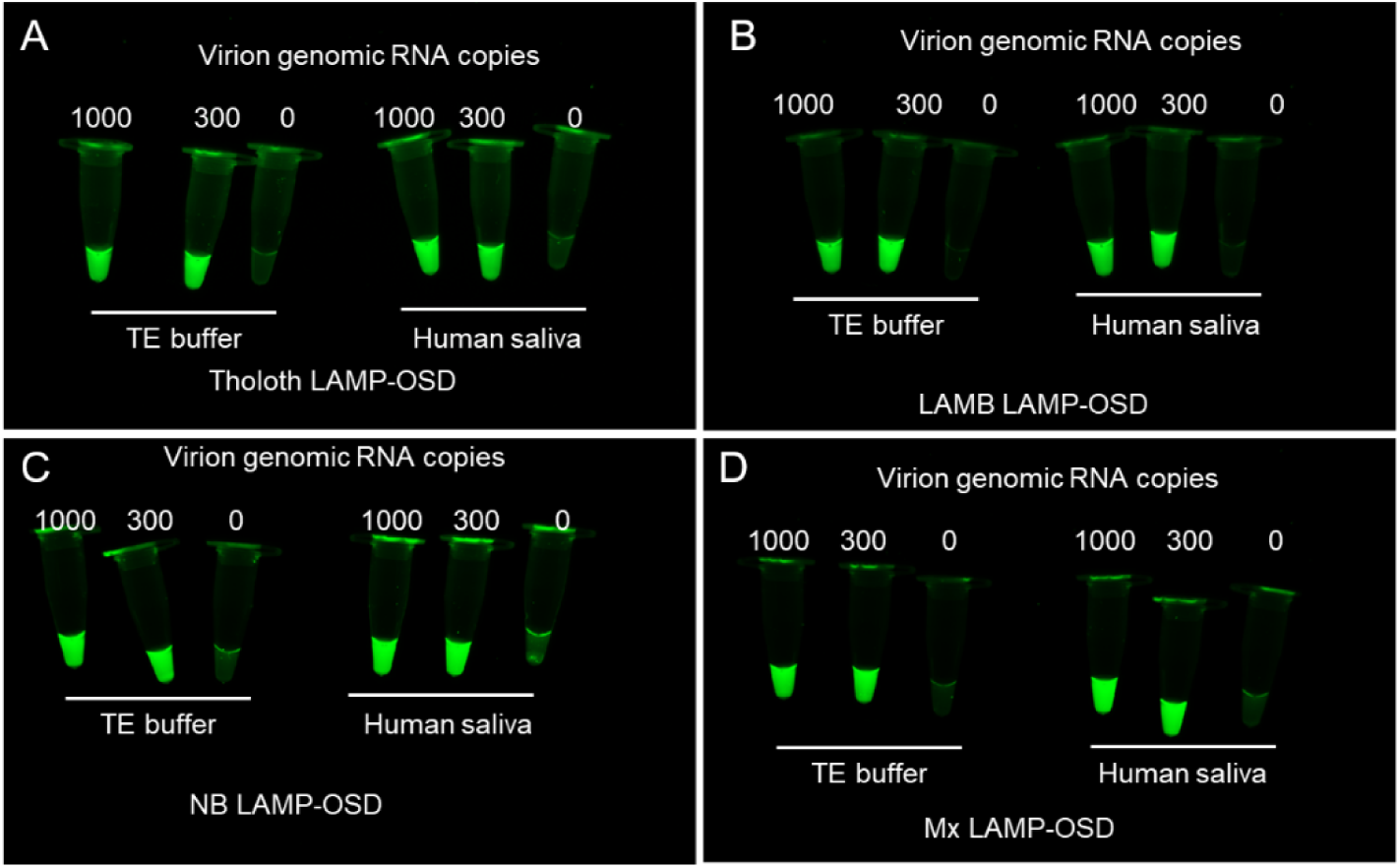
LAMP-OSD analysis of human saliva containing SARS-CoV-2 virions. Indicated virion amounts were spiked in TE buffer or human saliva and added to individual or multiplex (Mx) LAMP-OSD assays. Endpoint images of OSD fluorescence are depicted for Tholoth (A), 6 primer LAMB (B), and NB (C) individual LAMP-OSD assays and Tholoth+NB Mx LAMP-OSD assays (D).

### SARS-CoV-2 LAMP-OSD using Bst-LF

Although there are several DNA polymerases (such as Bsu, Phi29, Klenow, Vent, and Deep Vent) that can catalyze strand displacement DNA synthesis, Bst DNA polymerase has been reported to be the most efficient at performing LAMP.^*1*^ For the detection of RNA targets, reverse transcription LAMP commonly uses a combination of Bst and a dedicated reverse transcriptase, such as AMV reverse transcriptase. That said, it is known that some targets and primers are amenable to reverse transcription LAMP using only the Bst enzyme. The Bst DNA polymerase large fragment (Bst-LF; the precursor to the commercial version Bst 2.0 from New England Biolabs) possesses innate reverse transcription ability and can be used for LAMP.^*1*, *26*^ Moreover, Bst-LF can support LAMP even when used in the form of cellular reagents, dried bacteria containing overexpressed enzymes that can be added directly to reactions (see also below).^*20*^

Considering potential supply chain issues, especially for resource poor settings, we sought to determine whether Bst-LF (an open source enzyme that maybe produced on site, see also attached protocol for production, **Supplementary information**) as a cellular reagent could be used for LAMP-OSD assays with SARS-CoV-2 targets. The previously described NB, Tholoth, and 6 primer Lamb LAMP-OSD assays were carried out with SARS-CoV-2 viral genomic RNA and Bst-LF enzyme as a cellular reagent^*20*^ in an *E. coli* BL21(DE3) derivative ΔendA ΔrecA strain (see also attached protocol for production, **Methods**).

Following 90 min incubation at 65 °C, visual observation of endpoint OSD fluorescence revealed that the Tholoth LAMP-OSD assay containing 1000 copies of SARS-CoV-2 genomic RNA was brighter compared to assays containing fewer or no SARS-CoV-2 genomic RNA (**Figure 7**). Similarly, 6 primer Lamb LAMP-OSD assays containing as few as 100 SARS-CoV-2 genomes were bright compared to controls containing no viral genomes. In contrast, all tubes of NB LAMP-OSD assay demonstrated background fluorescence with no virus-specific increase in OSD signal. These results are consistent with previous reports from NEB, the manufacturer of Bst 2.0 (https://www.neb.com/products/m0374-bst-3-0-dna-polymerase#Product%20Information), that while Bst-LF cannot support all reverse transcription LAMP reactions, for the Tholoth and Lamb primer sets it is suitable for the detection of SARS-CoV-2.

**Figure 7.**
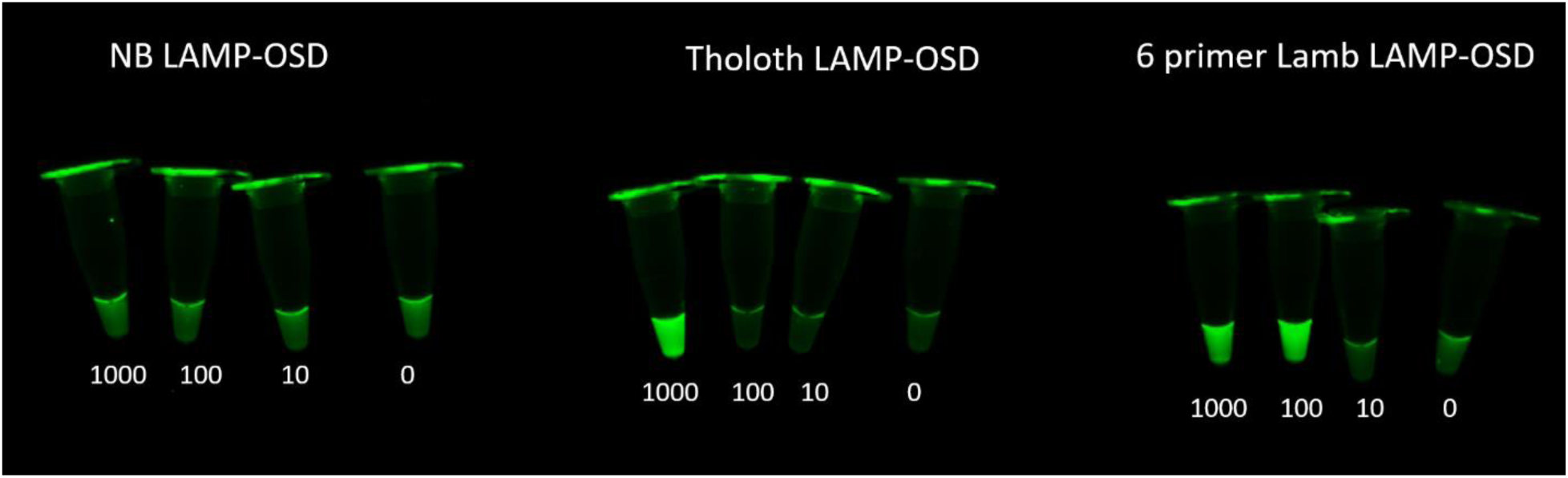
Execution of SARS-CoV-2 LAMP-OSD assays using only Bst-LF. Copies of SARS-CoV-2 genomic RNA used in each reaction are indicated below each tube. Image of OSD fluorescence at amplification endpoint is depicted.

### Conclusions and Resources

In summary, we have demonstrated a facile way to rapidly configure LAMP assays for accurate probe-based readout of SARS-CoV-2 by integrating OSD probes into individual and multiplex assays. The SARS-CoV-2 LAMP-OSD assays can be executed in one-pot reactions assembled using individual reverse transcription LAMP reagents, including with trivially produced cellular reagents. Moreover, these assays can be used to directly analyze crude human samples, such as saliva, to detect SARS-CoV-2 without interference from spurious signal. A few hundreds to a few tens of virion genomic RNA can be identified using individual or multiplex LAMP-OSD assays, respectively. We suggest that while LAMP-OSD may not have the same sensitivity as ‘gold standard’ RT-qPCR assays, that the versatility of LAMP-OSD, especially for resource poor settings with limited infrastructure, might prove useful for screening for positives, which could then be followed up with more limited or difficult to execute RT-qPCR tests.

Bst-LF purification is described in Reference *27* and detailed as a protocol in **Supplementary information**. Expression plasmids will be available from Addgene shortly, or can be obtained via https://reclone.org/. The production of cellular reagents is described in Reference 20 and detailed in the **Methods** section.

## Acknowledgements

We acknowledge Dr. Andre Maranhao for kindly providing a protocol for expression and purification of Bst-LF enzyme. We also acknowledge the FRI DIY Spring 2020 Peer Mentor Cohort for their project initiation and screenings of the preprint COVID-19 primer sets; Emma Aldred, Patricia Cadar, Elvi Casia, Haroon Dossani, Alijah James, Madi Kahanek, Andrew Kan, Shivani Kottur, Niam Kuttanna, Rory Malek, Maggie Miller, Morgan Motakef, Vylan Nguyen, Maria Nguyen, Nicole Nyamongo, Samantha Outar-Sankarpersad, Purvee Patel, Shiv Patel, Kush Patel, Lisa Sanchez, Sophia Solomon, Malaika Tah, Jose Torres, Reshan Warnesuriye, Sofia Williams, Eric Yang, and Julia Zheng. The Freshman Research Initiative (FRI)^33,34^ DIY Diagnostics Research Lab^35^ at the University of Texas at Austin develops point-of-care diagnostics with undergraduate students.

## Supplementary information

### Purification of Bst-LF or equivalent enzyme

For sites wishing to produce their own LAMP enzymes, the following protocol can be used.

|  |  |
| --- | --- |
| **Lysis Buffer** | • 20 mM Tris, pH 7.4<br>• 300 mM NaCl<br>• 0.1% Tween-20 (Thermo Fisher Scientific)<br>• 10 mM imidazole<br>+<br>• 1x EDTA-free protease inhibitor tablet<br>+<br>• 0.5 mg/mL HEW Lysozyme |
| **Wash Buffer** | • 20 mM Tris, pH 7.4<br>• 300 mM NaCl<br>• 0.1% Tween-20 (Thermo Fisher Scientific)<br>• 40 mM imidazole |
| **Elution Buffer** | • 20 mM Tris, pH 7.4<br>• 300 mM NaCl<br>• 0.1% Tween-20 (Thermo Fisher Scientific)<br>• 250 mM imidazole |
| **Dialysis Buffer** | • 50% Glycerol<br>• 10 mM Tris, pH 7.4<br>• 100 mM KCl<br>• 1 mM DTT<br>• 0.1 mM EDTA<br>• 0.5% Tween-20<br>• 0.5% Triton-X100 |

In brief, an *E. coli* protein expression strain such as BL21 was transformed with Bst-LF expression plasmid that contained an amino terminal six histidine tag for purification. An individual transformant colony was picked into desired media with appropriate antibiotics (e.g. ampicillin) and 1-2% glucose. This starter culture was grown overnight in shaking incubator at 37°C.

On day three, cultures were diluted 1:200 or 250 mL to 1 L of fresh medium. Inoculated cultures were grown to OD_600_ between 0.5-1 before inducing protein expression by adding anhydrotetracycline (aTC) to a final concentration of 200 ng/mL aTC. Induced cultures were allowed to express 3-7 hr for expression. Following expression, cells were harvested by centrifuging cultures at 4°C, 5000 xg for 20 min and then flash frozen in liquid nitrogen. Flash-frozen cell pellets may be stored at −80°C or allowed to thaw on ice for same day purification.

After thawing on ice, resuspend cell pellets in 30 mL cold **Lysis Buffer**. Transfer the resuspended cell pellet to a small 50 mL beaker with clean stir bar and place securely in an ice bath. With moderate stirring, sonicate the sample using 40% amplitude and 1 sec ON / 4 sec OFF for 4 min total sonication time.

Following sonication, centrifuge the resulting lysate at 4°C, 35,000 xg for 30 min. Carefully transfer the supernatant to a clean ultracentrifugation tube and incubate in a thermomixer at 400 rpm, 65°C for 20 min. Then place the heat-treated lysate on ice for 10 min. Centrifuge the heat-treated lysate at 4°C, 35,000 xg for 30 min. Carefully transfer the supernatant to a clean tube and filter the clarified lysate using a 0.2 μm filter.

Add clarified, filtered lysate to 1 mL of pre-equilibrated HisPur Ni-NTA resin and incubate for 30 min at 4 °C with end-over-end mixing for batch binding. Apply lysate and resin slurries to gravity columns and allow to drain. Wash columns three times with 10 mL of **Wash Buffer** and then elute with four 1 mL aliquots of **Elution Buffer**. Finally, pool elutions and dialyze overnight into **Dialysis Buffer** and store at −20 °C.

